# Twist, Snail, and Sox9 form an allosterically regulated complex, the EMTosome, on a bipartite E-box site

**DOI:** 10.1101/2020.02.03.932962

**Authors:** Daniel S McCracken, Hongzhuang Peng, Kasirajan Ayyanathan, Yike Lindy Jiang, Sarah Welsh, Jing Yang, Eileen Kennedy, Frank J. Rauscher, Alessandro Gardini, Frank J Rauscher

**Affiliations:** Biochemistry and Molecular Biophysics Graduate Group, Perelman School of Medicine, University of Pennsylvania, 3400 Civic Center Boulevard, 19104, Philadelphia, PA, U.S; The Wistar Institute, Philadelphia, Pennsylvania 19104, USA; Department of Pharmacology, Moores Cancer Center, University of California at San Diego, La Jolla, California, 92093, USA; Department of Pediatrics, University of California at San Diego, La Jolla, California, 92093, USA; Department of Pharmaceutical and Biomedical Sciences, College of Pharmacy, University of Georgia, Athens, GA 30602

**Author notes:** Contact Information, Corresponding Author: Frank J Rauscher III, 3601 Spruce Street Philadelphia, PA 19104. **Author contributions** Conceptualization was done by DSM, HP, FJRIII. Data Curation, Formal Analysis, Validation, Visualization were done by DSM, HP. Resources were done by DSM, HP, YLJ, FJRIV. Funding Acquisition was done by JY, FJRIII. Investigation was done by DSM, HP. Methodology was done by DSM, HP, KA, SW, JY, FJRIII, FJRIV. Project Administration was done by FJRII. Supervision was done by HP, AG, FJRIII. Writing – Original Draft Preparation was done by DSM. Writing – Review and Editing was done by all authors.

**Keywords:** transcription factors, epithelial mesenchymal transition, cancer, metastasis, gene regulation, EMTosome, Twist, Snail, Sox9

## Abstract

Epithelial-Mesenchymal transition (EMT) of primary tumor cells is a critical trans-differentiation event that contributes to dissemination and metastasis. The process of EMT is controlled by specific DNA-binding transcription factors (TFs) that reprogram the tumor transcriptome. In particular, the canonical EMT-TFs Twist and Snail can induce an EMT program when overexpressed in cancer cells, and both are found upregulated in metastatic cancers. Twist and Snail bind DNA directly, by recognition to variants of the E-Box sequence CANNTG. However, it is unclear how this binding is regulated. We have used a biochemical approach to dissect DNA binding and protein-protein interactions that occur amongst these proteins. We find that Twist preferentially recognizes a dyad repeat of E-boxes that are not directly bound by Snail. Our data suggest that Twist use its WR domain to recruit Snail into a binding complex through the Snail zinc-finger motifs. We analyzed Twist-Snail complexes in the breast carcinoma cell line SUM1315 and found evidence that it contains an additional protein partner, Sox9. Notably, we report that a native Twist complex can be displaced from its dyad binding site by consensus DNA binding sites for Snail and Sox9 even though these proteins do not contact the Twist dyad site. Taken together, our findings suggest that Snail and Sox9 interact with Twist to regulate its DNA binding ability via protein-protein interactions, thereby allosterically regulating Twist DNA binding. We designate this ternary complex EMTosome. These results may inform efforts to therapeutically target the EMT program in order to target cancer metastasis.

## Introduction

Epithelial-Mesenchymal Transition (EMT) is a genetic regulatory program involved in the differentiation and dedifferentiation of cells which enables them to become motile (1). This process occurs during developmental differentiation in many tissues, notably the neural crest, and during wound healing. Cancer pathobiology often includes reactivation of this program for the purpose of metastasis, where it provides the motility necessary to leave the primary tumor, and enter the circulation for distant spread, as well as therapeutic resistance (2). EMT is regulated on a transcriptional level by a series of proteins, considered master regulators of EMT. The major proteins involved are Twist, Snail, Slug, and Zeb (3). Further understanding of EMT is critical to our understanding of cancer and metastasis, as these proteins have been shown to be indicators of poor outcomes in cancer treatment (4).

Twist is a transcriptional activator and a type II basic helix-loop-helix (bHLH) protein that requires an obligate binding partner in order to regulate transcription (5). This binding partner is typically E12 or E47, both encoded by *E2A*/*TCF3*, with which it forms a dimer of heterodimers, or a tetramer (6). These binding partners are type I bHLH proteins, and all are part of the HLH protein superfamily (5). We have previously discovered that the optimal Twist binding site occupied in cells to is a dyad E-box (<u>CANNTG</u>NNNNN<u>CANNTG</u>) binding motif spaced by 5 nucleotides, such as the one in the *Alpha-2-macroglobulin* (*A2M*) gene, in order to regulate transcription (6). The WR domain of Twist dimerizes Twist, thereby forming a tetramer of Twist-E47 molecules on the dyad site. The WR domain also serves as a platform for protein-protein interactions (PPIs) which occur because of the tetramer binding DNA. Snail is a transcriptional repressor and a zinc finger protein, also a master regulator of EMT, that also binds to subsets of E-box sites which appear to be different than those for Twist: the specificity to differentiate between E-boxes is determined by the central NN nucleotides (CA<u>NN</u>TG) (6) and partially by flanking sequences. A classic, canonical DNA binding site for Snail is located in the *E-cadherin* gene (*E-cad*) (6). The role of Snail in metastasis, tumor progression, and endowing a stem-like phenotype as part of its role in EMT and related processes has been well documented (3, 7, 8) and has broad impacts on tumor growth and invasiveness (9). Sox9 is a transcriptional regulator which recognizes a specific DNA sequence (GACAATG) and is part of the High Mobility Group (HMG) family of proteins (10), and has been directly implicated in EMT during both development and cancer progression (11, 12). Twist has been shown to inhibit the transcriptional regulatory functions of Sox9 (13), and it may impact the stabilization of Snail as well (14). However, how these proteins function together to regulate transcription of genes involved in EMT, given their various functions has been unclear. We sought to use biochemical methods to dissect an endogenous complex to clarify the manner of DNA binding and transcriptional regulation by these proteins.

## Materials and Methods

### Cloning and Vectors

All cloning was done by PCR amplification and subcloned into the respective plasmids. The pCDNA3.1-Sox9-C-Flag was acquired from Genescript and used as the template for cloning GST-Sox9/DIM+HMG (Forward primer 5’ – *ATACGCGGATCCAATCTCCTGGACCCCTTCAT* – 3’ and reverse primer 5’ – *ATTCCGGGATTCTCAGATGTGCGTCTGCTCCGTG* – 3’) into pGEX4T1 plasmid and His-Sox9/DIM+HMG (Forward primer 5’ – *CGCGGATCCAATCTCCTGGACCCCTTCATG* – 3’ and reverse primer 5’ – *CCCAAGCTTTCAAGGTCGAGTGAGCTGTGT* – 3’) into pQE30 plasmid. pCDNA3.1-Snail-C-Flag and pCDNA3.1-HA-Snail were from our laboratory and used as a template for cloning His-SnailZF1-4 (Forward primer 5’ – *TCGCACACGTGCCTTGTGTCTGCA* – 3’ and reverse primer 5’ – *CTGCTAAGTCGCTCCCGGAGGCCT* – 3’) into pQE30 plasmid. pGEX4T1-SnailZF1-4 (GST-SnailZF1-4) was from our laboratory (described earlier). The pCDNA3.1-Twist and pQE-Twist were as previously described (15) and used as the template for cloning GST-WR (Forward primer 5’ - *ATCATAGGATCCAGCTGCAGCTATGTGGCTCAC* – 3’ and reverse primer 5’ - *ATGCTGAAGCTTTTAGTGAGATGCGCTCATGGAC* – 3’) into pGEX4T1 plasmid, dual expressed His-Twist/bHLH+WR (Forward primer 5’ - *ATCATAGGATCCACAGTCTTACGAGGAGCTCCAA* – 3’ and reverse primer 5’ - *ATGCTGAAGCTTTTAGTGAGATGCGCTCATGGAC* – 3’) into pETDuet plasmid, first cloning site, and His-Twist/bHLHΔWR (Forward primer 5’ - *ATCATAGGATCCACAGTCTTACGAGGAGCTCCAA* – 3’ and reverse primer 5’ - *ATGCTGAAGCTTCTAATCCAGCTCATCAGACTGCAG* – 3’) into pETDuet plasmid, first cloning site. The pCDNA3.1-E47/bHLH was as previously stated (15) and used as the template for cloning dual expressed E47/bHLH (Forward primer 5’ - *ATCATAAGATCTCTGTCCCTGGAGGAGAAAGAC* – 3’ and reverse primer 5’ - *ATGCTGCTCGAGCTAATTCAGGTTCCGCTCTCGCAC* – 3’) into pETDuet plasmid, second cloning site.

### Cell Culture and transfections

Sum1315 cells were acquired from the Zhou lab at the University of Kentucky, and originally developed by Rosenblatt lab (16) and were cultured in F-12 media (Gibco 11765-047) supplemented with 5% FBS (Corning MT35-010-CV), 2mM Glutamine (Corning 25-005-CI), 50units/mL Pen Strep (Gibco 15140-122), 10 ng/mL EGF (Corning 354001), 5 μg/mL Insulin (Gibco 51500-056). Cos-1 cells were acquired from ATCC and cultured in DMEM (Corning 10-017-CV) supplemented with 10% FBS (Corning MT35-010-CV), 2mM Glutamine (Corning 25-005-CI), and 50units/mL Pen Strep (Gibco 15140-122). Transfections were done with pCDNA3.1 recombinant plasmid. Prior to transfection, Cos-1 cells were grown to 80-90% confluency in 15cm tissue culture dishes and media was changed to Pen Strep free media 12-16 hours before transfection. Before transfection, media was changed to Opti-mem, with no additives. Transfections were done using Lipofectamine 2000 (Invitrogen 11668-019). For single transfections, 20μg of DNA was used. For triple transfections, 24μg DNA was used, 8μg of each plasmid. For quadruple transfections 24μg of DNA was used, 6μg of each plasmid. DNA being transfected was mixed in 1.5mL Opti-mem and incubated for 5 min at 25°C. 60uL of Lipofectamine was mixed with 1.5mL Opti-mem and incubated for 5 min at 25°C. DNA and Lipofectamine Opti-mem were then mixed and incubated for 30 minutes at 25°C. Then the Lipofectamine/DNA mixture was added to the cell cultures and incubated for 4 hours at 37°C in incubator. Media was changed back to Pen Strep free media and incubated for 48 hours. Nuclear extracts were then made utilizing the provided nuclear extract protocol.

### Nuclear Extracts and Whole Cell Lysates

For nuclear extracts, cells were cultured to 90% confluency in 15cm dishes. All steps were performed at 4°C. Cells were rinsed four times with 5mL PBS. 10 mL of Lysis Buffer (10mM HEPES, 10mM NaCl, 1.5mL MgCl_2_, 20% Glycerol, 0.2mM EDTA, 0.1% Triton X-100, 1mM DTT, 1mM PMSF, 10µg/μL pepstatin, 10μg/µL leupeptin, 10μg/µL aprotinin, 2mM Sodium Ortho-vanadate) was added and incubated for 2 minutes on ice. Cells were scraped and incubated 4 minutes on ice. Following that, cells were scraped into 50mL conical tubes and centrifuged at 500g for 10 minutes at 4°C. Supernatant was removed and pellets were resuspended in 1mL of Nuclear Extract Buffer (10mM HEPES, 500mM NaCl, 1.5mL MgCl_2_, 20% Glycerol, 0.2mM EDTA, 0.1% Triton X-100, 1mM DTT, 1mM PMSF, 10μg/μL pepstatin, 10μg/μL leupeptin, 10μg/μL aprotinin, 2mM Sodium Ortho-vanadate). Pellet was resuspended by gentle pipetting 60 times, incubated on ice 5 minutes, then gentle pipetting 40 times. The resuspended extract was rotated at 4°C for 90 minutes, followed by centrifugation at 13k RPM for 30 minutes. Extracts were then dialyzed into binding buffer (20mM HEPES, 100mM KCl, 1mM DTT, 10% Glycerol), aliquoted into 100μL portions and snap frozen. 40μ of nuclear extract was run on Western blot. Whole cell lysates were made by harvesting and lysing cells in a 10cm dish (80-90% confluency) in 100 μl of Tween 20 Lysis Buffer (17)(25 mM Tris/Hepes, pH 8.0, 25 mM NaCl, 2 mM EDTA, 1 mM PMSF, 0.5% Tween 20). Lysate was incubated on ice for 15 minutes and the volume was estimated. 1/5 volume of 5 x Buffer was added to lyse the nuclei. The final concentration of the lysis buffer is 1% Triton x100, 20 mM Tris, pH 8.0, 150 mM NaCl, 10 mM MgCl2, 2 mM EDTA, 10% Glycerol, 2.5 mM β-mercaptoethanol, 1 mM NaF, 1 mM DTT, 1 mM PMSF. Lysate was incubated on ice for 15 min, then centrifuged at 13k RPM for 10 min and the supernatant was recovered. 5 μl of supernatant was removed to check protein concentration by Bradford assay and 100 μg of whole cell lysate was used to do Western blot.

### Protein Expression and purification

For native purification, pETDuet-Twist-E47 plasmids were transformed into Rosetta2(DE3)-pLysS (Millipore) bacteria. Colonies were selected and grown in 2YT media 24 hours, and then used to start fresh cultures in 800mL of 2YT at OD_600_ of 0.1-0.2. These cultures were grown to OD_600_ 0.6-0.8 then protein production was induced with 500μM isopropyl-1-thio-β-D-galactopyranoside (IPTG) and induction was done 12-16 hours at 20C. Cultures were harvested by centrifugation and resuspended in 40mL of sonication buffer (50mM sodium phosphate pH 8.0, 500mMNaCl, 5 mM imidazole, 1mM PMSF, 500μM TCEP, 0.5% Tween-20, 10μg/mL DNaseI, 150μg/mL Lysozyme). Supernatant was sonicated for 3 minutes, 3x on ice (Branson Sonifier 450, 80% power, 100% duty) and centrifuged 12k RPM for 30 minutes at 4°C to clarify. Supernatant was bound to 500μL TALON Metal Affinity Resin (Takara 635501) pre-equilibrated in sonication buffer, by rotating at 4°C for 1 hour. Beads were then washed 4x with 10mL Wash buffer (50mM sodium phosphate pH 8.0, 500mMNaCl, 20 mM imidazole, 1mM PMSF, 500μM TCEP, 0.5% Tween-20). Protein was eluted from beads with elution buffer (50mM sodium phosphate pH 8.0, 500mMNaCl, 1M Imidazole). Eluted protein was then dialyzed in 4L SEC buffer (25mM HEPES pH 7.5, 200mM NaCl, 500μM TCEP) twice, 2 hours each, and then once 12-16 hours, all at 4°C. Protein was then concentrated to ∼20mg/mL using centrifugal concentrators (Amicon Ultra-15 3k NMWL UFC900324) and run on FPLC (AKTA Pure) for size exclusion chromatography (SEC column used is Superdex 75) and 1mL fractions were collected. After analyzing all fractions by SDS-Page, fractions 10 and 11 were collected and pooled as purified proteins. Expression of pQE plasmids, denaturing purifications, and refolding were done as previously described (15). GST-Fusion proteins were purified as previously described (18).

### Electrophoretic motility shift assays (EMSA)

Protocol for EMSAs, oligonucleotide annealing, probe labeling, and probes used are previously described (6), except as follows. The *A2M* DNA probe used is comprising the −2512 to −2531 promoter region of the human *A2M* gene, with a CG added to each end to provide additional stability. The 5’ to 3’ sequence is 5’ - *GCT<u>CAGATG</u>GAAGC<u>CATGTG</u>TGC* - 3’, with the binding regions, E-boxes, underlined. The Mut1 and Mut2 mutations are as previously described (6). The double mutant (Dbl Mut) probe was comprised of both E-box mutations. Antibody shifts were done by incubating the protein/extract with 4uL of antibody (α-Twist: Invitrogen PA5-47824, α-Snail: Santa Cruz E-18, α-Sox9: Abcam ab3667, α-E47: BD Pharmingen 554077) for 15 minutes at 25°C before adding radiolabeled probe. Equilibrium competition was done by mixing labeled and unlabeled competitor together and incubating for 15 minutes at 25°C prior to addition of extract/proteins. Order of addition competition was done by adding the unlabeled competitor after binding of protein to DNA, and then incubating for 30 minutes at 30°C. The competitor probes were as follows, 5’ to 3’, with the binding regions underlined, E-cad 5’-*GTGGCCGG<u>CAGGTG</u>AACCCTCA* - 3’, Sox9 5’-*CCAG<u>GACAATG</u>CCGC<u>CATTGTC</u>CTG*G - 3’, SZF 5’ – *GATCCCAGGGTAACAGCCGTTTG* – 3’.

### Immunoprecipitation and Western Blot

Immunoprecipitations and Western Blots were done as previously described except as noted below (19). Addition of oligonucleotides to NE increased affinity to flag antibody resin. 5μL of oligonucleotide (2mg/mL) and 10uL of dI: dC (1μg/μL) was added to each 1mL of nuclear extract, and then incubated at 30°C for 30min at 120 rpm. Affinity beads [Anti-Flag M2 affinity gel (Sigma A2220), Anti-Myc (Pierce 20169), Anti-HA (ICL RGHT-145D)] was pre-washed in BB buffer (20mM HEPES, 100mM KCl, 1mM DTT, 10% Glycerol) twice, and 1mL of nuclear extract was bound to 200μL of 50/50 slurry by rotating for 2 hours at 4°C. Resin was then washed five times with BB buffer. Protein was eluted twice from resin using 100μL of 500ng/μL [3x flag peptide (Sigma F4799), Myc (Pierce 20170), HA (Thermo Scientific 26184), respectively] by incubating at 37°C for 20 minutes shaking at 150rpm, and then supernatant separated from beads using centrifuge columns (Pierce 89868). Primary antibodies used for Western Blots were α-Twist (Santa Cruz 2C1A), α-Snail (Abcam ab63371), α-Slug (Cell Signaling C19G7), α-E47 (BD Pharmingen 554077), α-E12/E47 (BD Pharmingen 554199), α-Vimentin (Cell Signaling 5G3F10). Primary antibodies used are at 1:1000 dilution unless specified. Secondary antibodies are HRP conjugated α-mouse (7076), α-rabbit (7074), or α-rat (7077) from Cell Signaling, used at 1:5000.

### GST binding assays, Biotinylated Affinity Chromatography

GST association assays were performed as previously described (Peng, JBC, 2000) except as noted below. Binding was done at 37°C instead of room temperature. Wash steps were performed with the buffers BB200 (200mM NaCl, 20mM Tris pH 7.5, 10% glycerol), BB500 (500mM NaCl, 20mM Tris pH 7.5, 10% glycerol), and RIPA (10mM Tris pH7.4, 150mM NaCl, 1% Triton X-100, 1% DOC, 0.1% SDS). Antibodies used were the same as for Western Blots.

For biotinylated affinity chromatography, the same DNA probes were used as in EMSA, except for biotin being attached to the 5’ end of the DNA oligonucleotide (−2531 in relation to transcription start site). Magnetic streptavidin beads (Invitrogen Dynabeads M-280 Streptavidin– 10μL bead volume) were equilibrated in binding buffer previously described under nuclear extracts, by rotating for 5 minutes at 25°C. Supernatant was removed and binding buffer was added to a volume of 1mL, and biotinylated DNA (30μL of 20μM) and BSA (5μL of 200mg/mL) were added, then rotated for 1 hour at 25°C. During this time, extracts were incubated with dI: dC (4μL of 1μg/μL in 400μL of nuclear extract) and incubated for 30 minutes at 30°C. Supernatant for DNA binding to beads was removed, 600uL of binding buffer and extracts were then added to beads and incubated by rotating for 1 hour at 25°C. Beads were washed four times with binding buffer plus 0.2% NP40, rotating 5 minutes at 25°C for each. Beads were resuspended in 50uL of 1x Laemmli buffer and visualized by SDS-PAGE and Western Blot (α-Twist 2C1A, α-Snail L70G2, α-Sox9 Abcam ab3679).

### Chromatin Immunoprecipitation

COS-1 Cells were grown to 90% confluency in 15cm dishes and transfected as described in this manuscript. Cells were fixed with 1% Formaldehyde for 5 minutes at 25°C, then quenched with 125mM glycine for 5 minutes at 25°C. Cells were harvested and the chromatin was enriched in chromatin enrichment buffer (20mM Tris HCl (pH 8.0), 0.25% Triton X-100, 200mM NaCl, 10mM EDTA, 0.5mM EGTA, 1ug/mL Protease inhibitors [aprotinin, leupeptin, pepstatin], 1mM Benzamidine, 50mM NaF, 10mM NaOV3, 5mM butyric acid) for 10 minutes at 4°C. Cells were pelleted (5.0×10^6^ cells per 1mL) and resuspended in IP Buffer (20mM Tris pH 8.0, 0.05% DOC, 0.5% Triton X-100, 0.5% NP-40, 200mM NaCl, 5mM butyrate, and Protease Inhibitors), then sonicated to a peak of 200-500bp (Branson Sonifier 450, in ice bath, 80% power, 30 seconds sonication, 30 seconds incubating on ice, 10 minutes total sonication time). Chromatin was precleared with Protein A/ beads (20uL per 1mL), which were pre-blocked with salmon sperm DNA and BSA, for 1 hour at 25°C. 1mL of Chromatin was incubated 12-16 hours at 4°C with 10μg of antibody (α-RNA Pol2 [sera from Gardini lab], α-Twist [Abcam ab50887], α-Snail [R&D Systems AF3639], α-Sox9 [Abcam ab3697], Normal mouse IgG [Santa Cruz sc-2025]). Immune complexed were bound to 20uL pre-blocked Protein A/G Beads at 4°C for 3 hours. Washes were: 4x with IP Buffer, 2x with High Salt IP Buffer (400mM NaCl), 1x with LiCl buffer (10mM Tris pH 8.0, 250mM LiCl, 0.5% NP40, 1% Triton X-100, 0.1% DOC, 5mM EDTA, and PIs), 1x with Mixed Mycelle wash buffer (20mM Tris pH 8.1, 150mM NaCl, 5mM EDTA, 5% Sucrose, 1% Triton X-100, 0.2% SDS), 2x with TE. Elution was done with 250μL 50mM Tris (pH 8.0), 200mM NaCl, 5mM EDTA, 1% SDS on thermomixer at 37C for 30 minutes, twice. Crosslinks were reversed 12-16 hours at 65°C, then treated with 30ug of Proteinase K (Roche) at 55°C for two hours. DNA was purified by phenol/chloroform extraction and analyzed by standard PCR. *A2M* PCR Primers are: forward probe 5’-*CTCTGTCACATAAAGGTCCC*-3’, reverse probe 5’-*AAGGGTACAGTTCTGCGCTT*-3’. *E-cad* PCR Primers are: forward probe 5’-*TAGAGGGTCACCGCGTCTAT*-3’, reverse probe 5’-*TCACAGGTGCTTTGCAGTTC*-3’.

## Results

### Characterization of a Twist-protein complex in breast cancer cells

To begin to assess the protein-DNA complexes associated with Twist, we performed electrophoretic mobility shift assays using a canonical Twist dyad binding site from the *A2M* promoter, using nuclear extracts from the breast carcinoma cell line Sum1315, which is well-characterized for studying EMT (16). Additionally, we supplemented these analyses using purified proteins for Twist, Snail, E47 and Sox9 and derivatives, as well as their consensus DNA binding sites, as depicted in Figure 1A-C. Whole cell lysates and nuclear extracts from Sum1315 cells were confirmed to contain Twist, E47, Snail, and Sox9, as well as the other EMT markers, Slug and Vimentin (Sup Fig 1B). We utilized EMSA with ^32^P radiolabeled *A2M* DNA probe, and antibodies (Sup Fig 1C) to verify the existence of a complex and verify the presence of these proteins in that complex (Fig 1D). Nuclear extracts from Sum1315 cells produced a consistent and reproducible complex with the A2M DNA probe (Lane 2). This primary complex shows altered mobility when Twist antibody is added (Fig 1D, Lane 3), verifying the presence of Twist protein within this complex as expected. Surprisingly, the integrity of this primary complex is greatly reduced by addition of Snail or Sox9 antibodies suggesting that both Snail and Sox9 are additional binding partners in this multiprotein complex. (Fig 1D, lanes 4 and 5). For clarity we denote this protein-DNA complex the “EMTosome”.

**Figure 1.**
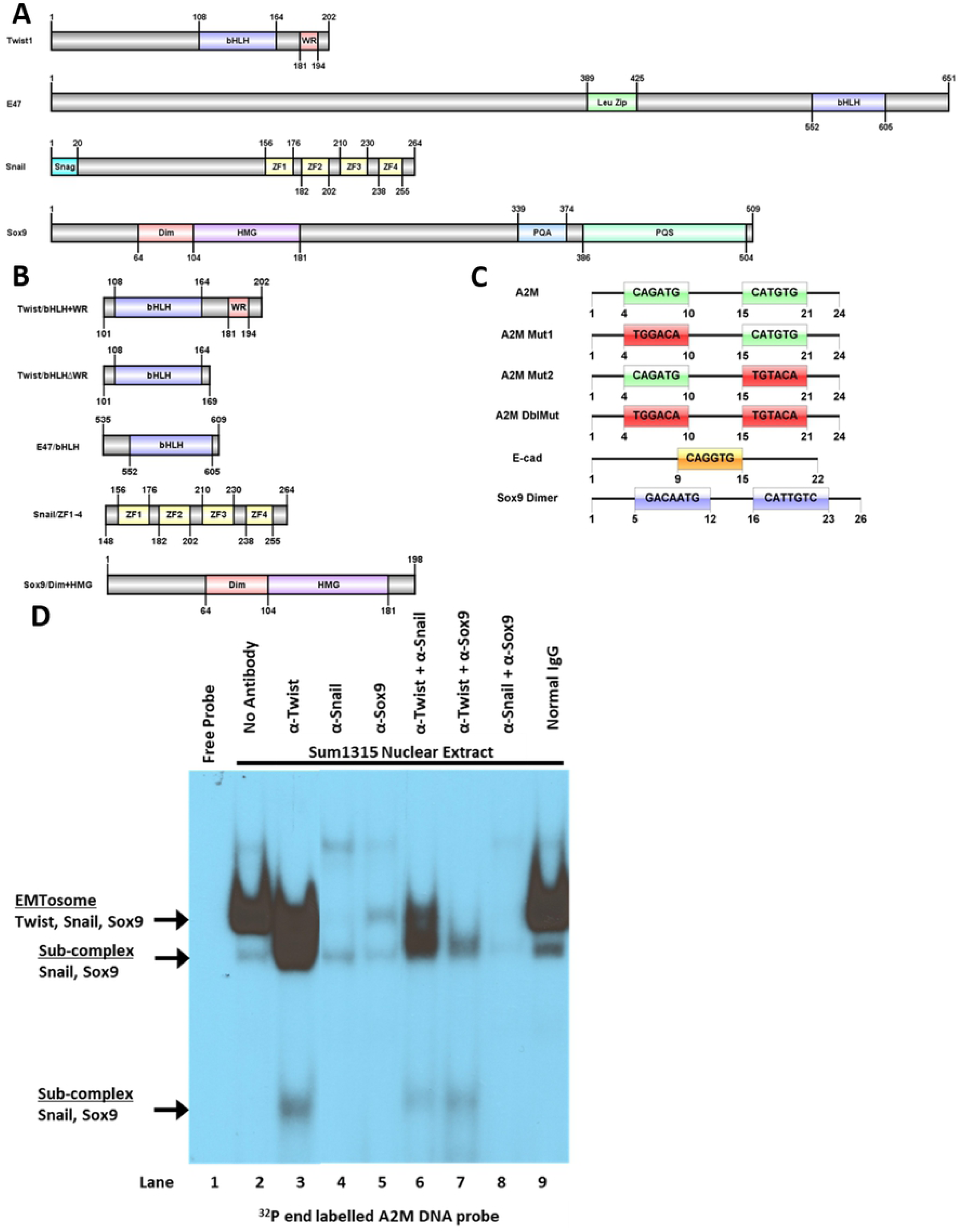
The EMTosome complex contains Twist, E47, Snail, and Sox9, which all interact with different consensus DNA sequences. A) Diagrams of full-length Twist, E47, Snail, and Sox9. B) Diagrams of recombinant proteins used for this study. C) Diagrams of oligonucleotides used in our study, showing the double E-box of the *A2M* oligo, the mutations used to eliminate E-box specificity of binding, the single E-box of E-cad binding site, and the consensus DNA binding site of a Sox9 dimer. D) Electrophoresis Motility Shift Assay (EMSA) utilizing radiolabeled *A2M* DNA probe utilizing antibodies to create motility shifts, showing the responsiveness of the EMTosome to antibodies to Twist, Snail, and Sox9.

We next supplemented these analyses using purified proteins from *E. coli* such that their native DNA binding specificities could be ascertained in the absence of associated proteins or post-translational modifications. The full-length Snail and Sox9 are unstable in *E.coli*, however removing the N-terminal region from Snail and the C-terminal region from Sox9 greatly stabilized them. After purification with a 6xHis tag utilizing a denaturing/renaturation protocol they can be prepared as highly soluble well-folded proteins (Sup Fig 2A). We found early on that Twist and E47 produced in *E. coli* are only stable if co-expressed. To achieve this, Twist and E47 sequences, with codons optimized for bacterial expression were cloned together into a dual expression vector (pETDuet) and purified under native conditions for cobalt chromatography, then further purified with size exclusion chromatography (SEC). The protein complex was verified by Coomassie staining and was confirmed by Western Blot (Sup Fig 2B). Fractions 10 and 11 from the SEC were used for further assays. The sizes of the recombinant proteins are 13kDa (Twist/bHLH+WR) and 9kDa (E47/bHLH), but they elute from the column at ∼44 kDa and larger, suggesting either extended, non-globular secondary structure, or tetramer formation. Using these highly pure proteins we tested the ability of Snail to bind to Twist and Sox9. Utilizing a GST-Snail-ZF1-4 fusion protein supplemented with nuclear extracts from Sum1315 cells, we detected associations between Snail-ZF1-4 and both Twist and Sox9, reproducibly and under harsh washing conditions (Fig 2A). The GST-Twist-WR domain shows association with Twist and Snail from lysates, suggesting that the WR domain of Twist is necessary for interactions with endogenous Twist and Snail (Fig 2A right panel). Notably, a GST-Sox9/DIM+HMG fusion protein was also able to associate with Snail/ZF1-4 recombinant protein, providing evidence that the interaction between Snail and Sox9 may be direct (Fig 2B). These combined data support an interaction between Snail and both Twist and Sox9 through the zinc finger region of Snail, and an interaction between Twist and Snail through the WR domain of Twist, supporting our observations of the native EMTosome in SUM1315 Cells.

**Figure 2.**
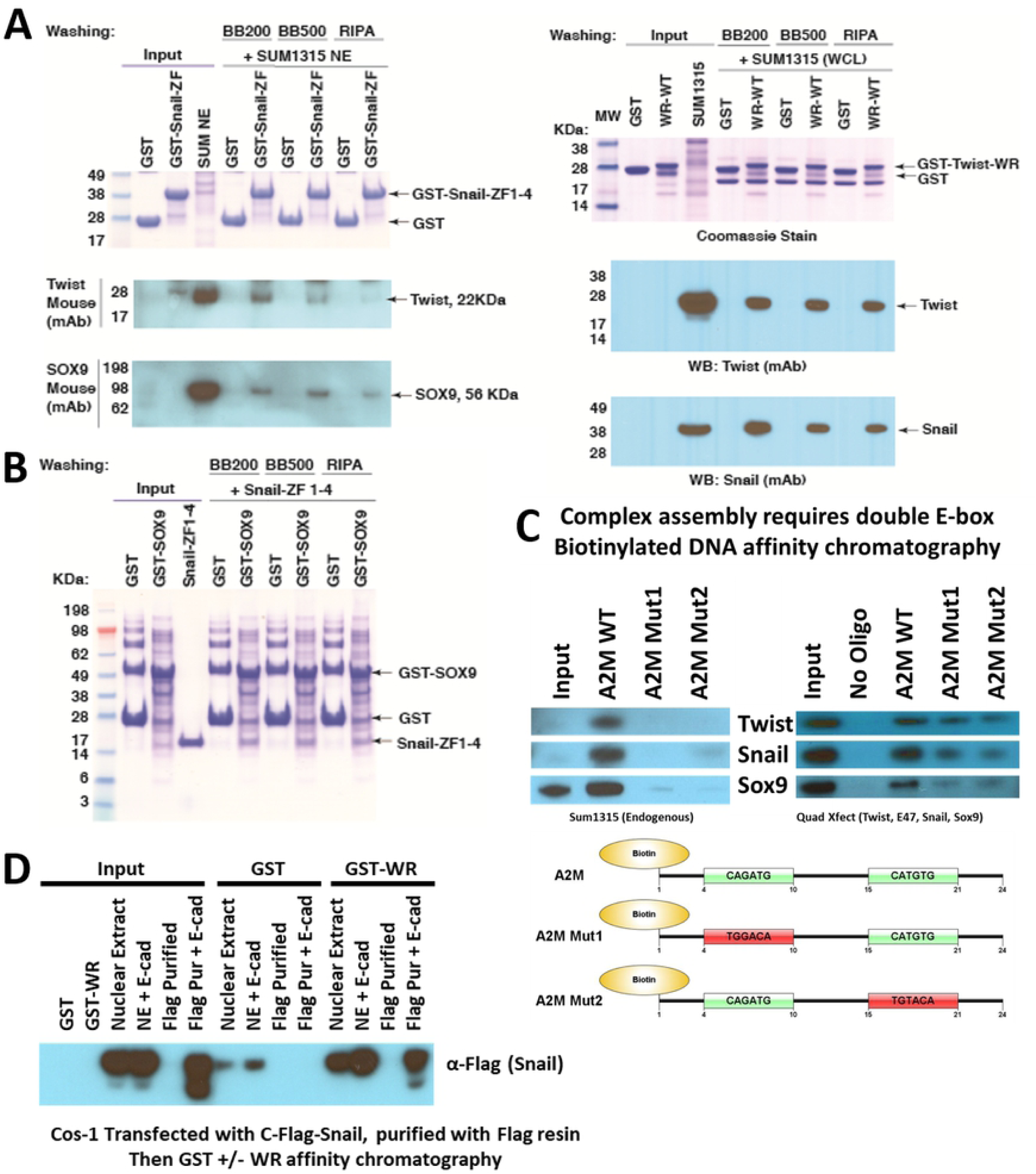
All members of the EMTosome protein complex interact with each other through direct or indirect interactions in a variety of conditions and can be shown to copurify via affinity chromatography. A) GST-Snail/ZF1-4 affinity chromatography from Sum1315 nuclear extracts and GST-WR affinity chromatography from Sum1315 cell lysates with both coomassie staining and Western Blot shows that the Snail ZF region is capable of binding with Twist and Sox9 and the Twist WR domain is capable of binding with Twist and Snail. B) GST-Sox9 affinity chromatography with recombinant purified Snail/ZF1-4 with coomassie staining shows that interaction between Sox9 and Snail is direct. C) Western Blot of affinity chromatography from both endogenous (Sum1315) and transfected (Cos-1; Twist, E47, Snail, and Sox9 transfected) nuclear extracts using biotinylated DNA probes of the A2M WT binding site and mutants shows that the dyad E-box has much greater affinity for the EMTosome complex than either individual E-boxes. D) Western Blot of flag epitope purification of Snail-C-Flag from transfected Cos-1 nuclear extracts and then GST-WR affinity chromatography shows that the interaction between Twist and Snail is direct with the Twist WR domain.

We next analyzed the impact of each E-box DNA site contained in the dyad target on the ability of the EMTosome to form using SUM1315 nuclear extracts. Utilizing biotinylated *A2M* DNA binding sites, we found that the dyad E-box is required, as mutating either of the E-box binding sites abolished the interaction between Twist, Snail and Sox9 in this complex (Fig 2C, left panel). Similar data were obtained in Cos-1 cells transfected with all four proteins (Fig 2C, right panel). We also found that the ability of the WR domain of Twist to interact with flag-purified Snail was facilitated by the addition of the DNA binding site for Snail from the *E-cadherin* promoter (E-cad, Fig. 2D). These data suggest that DNA binding by Snail derived from *E. coli* may help Snail fold into a confirmation suitable bind to the Twist WR domain, as well as show a direct interaction between Twist WR domain and Snail.

### DNA Binding characteristics of the EMTosome

To further characterize the DNA binding properties of the EMTosome, we analyzed the effects of competitor DNA (Fig 3A) on the endogenous complex in Sum1315 nuclear extracts using EMSA under both equilibrium and non-equilibrium conditions. By mixing radiolabeled *A2M* DNA binding site and an unlabeled probe (*A2M, E-cad* or Sox9) at a 20x concentration. Surprisingly, we observed that binding of the SUM1315 EMTosome is eliminated by any of these three sites (Fig 3A, Lane 3) when both hot and cold probes are mixed prior to protein addition. Alternatively, when the EMTosome is first bound to radiolabeled *A2M* DNA, even the homologous probe cold *A2M* was no longer capable of eliminating binding (Fig 3A, Lane 6), suggesting a low rate of dissociation of the complex once bound to DNA. Cold probes for *E-cad* and Sox9 binding sites, when added after EMTosome formation, also had modest ability to diminish complex formation (Fig 3A lanes 7 and 8).

**Figure 3.**
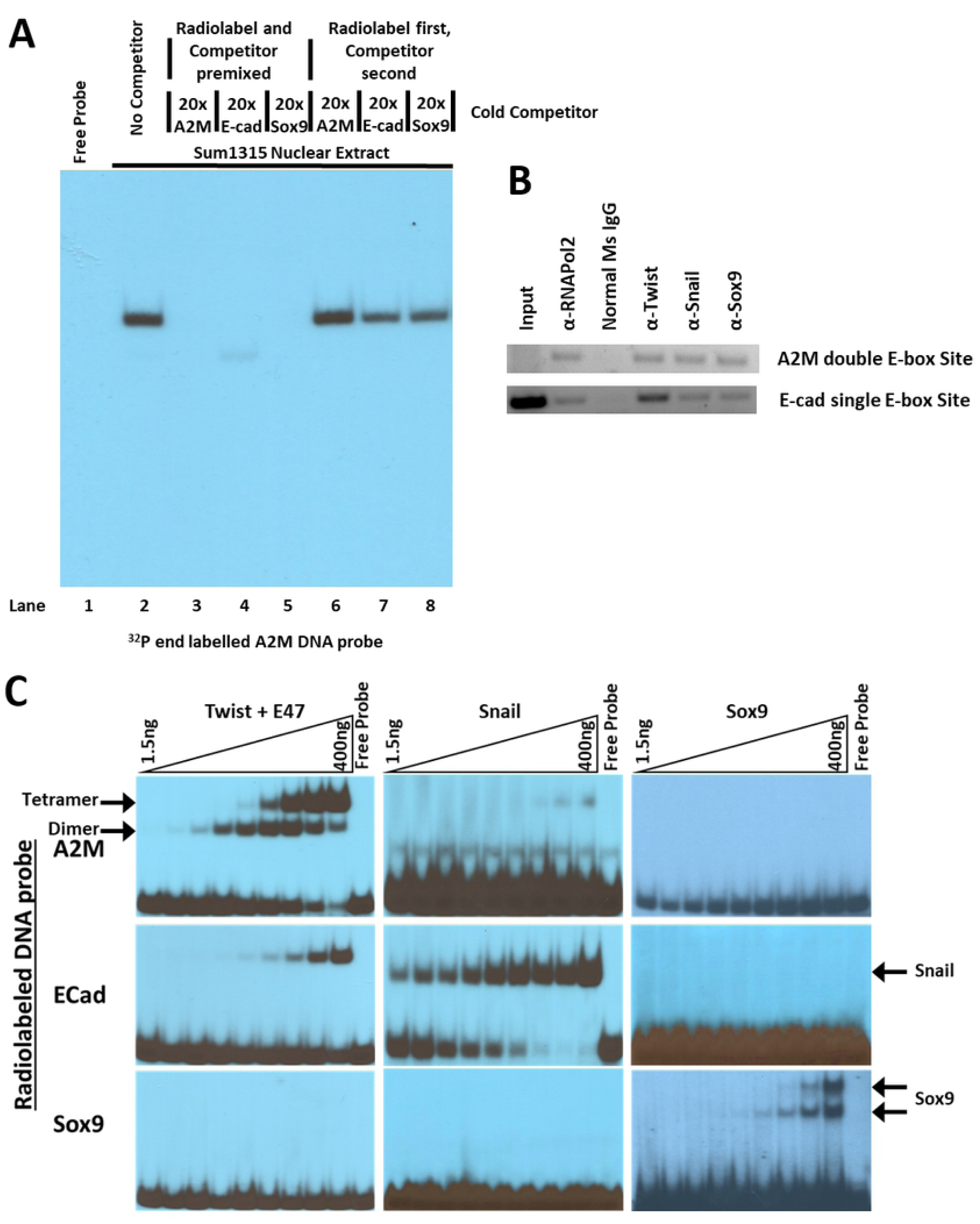
EMTosome binding to the A2M DNA target can be disrupted by the DNA targets of any of the members, but once the EMTosome is bound to the A2M DNA, it is largely refractory to cold competition. A) EMSA with radiolabeled A2M DNA probe with Sum1315 Nuclear Extract and competition with A2M, E-cad, and Sox9 unlabeled probes, both in equilibrium of hot and cold probes, and binding of extract to radiolabeled probe first. B) ChIP from Quadruple transfected Cos-1 Cells showing cohabitation of EMTosome proteins on A2M and E-cad transcriptional binding regions. C) EMSA of Twist/bHLH+WR, Snail/ZF1-4, and Sox9/DIM+HMG recombinant proteins titrated against A2M, E-cad, and Sox9 radiolabeled probes shows high specificity for each protein for their own consensus DNA sequences.

To confirm and extend these findings, we performed chromatin immunoprecipitation in cells expressing the core EMTosome components Twist, Snail, E47 and Sox9. We found that antibodies to Twist, Snail and Sox9 were all capable of immunoprecipitating a Twist canonical binding target, the dyad E-box site on the promoter region of the *A2M* gene, and a canonical Snail target, the single E-box site in the promoter region of the *E-cadherin* gene (Fig. 3B). These data support the ability of these three proteins to occupy the appropriate site in mammilian cells.

We next used purified recombinant proteins to determine the specificity of each protein for recognition of its canonical DNA binding site and for cross recognition amongst the EMTosome components. This detailed titration analysis revealed that each protein (or protein complex in the case of Twist and E47) showed strong specificity for their own canonical site but did not bind to sites recognized by the other components. (Fig. 3C). Of most importance, Snail showed very little (100-fold less) apparent affinity for the dyad *A2M* E-box twist sites. Similarly, the Twist-E47 complex bound very poorly to the canonical Snail site from *E-Cad*.

Neither Snail nor Twist-E47 proteins bound appreciably to the Sox9 site whereas Sox9 bound it robustly. Thus, the observation (Fig 3A) that E-cad and Sox9 sites can abolish binding of the EMTosome when bound to the dyad E-box cannot likely be accounted for by competition at the level of DNA binding specificity overlap amongst the probes. An alternative explanation is that Snail and Sox9 bind to Twist-E47 via protein-protein interactions only, and that occupancy of Snail and Sox9 with their canonical sites alters these protein-protein interactions with the Twist WR domain. We tested this by adding recombinant Snail and Sox9 to a complex of purified Twist-E47 bound to the dyad site. We observed the more slowly migrating band, previously documented as a tetramer formation of Twist and E47, increase in abundance (Fig 4A, Lanes 3 and 4 compared to 10 and 11). Notably, removal of the WR domain of Twist eliminates the ability of this higher order complex to form, which can be partially recovered through addition of Snail and Sox9 (Fig 4B). These data suggest that Snail and Sox9, while unable to bind directly to the A2M dyad site, can influence Twist-E47 binding to the site.

**Figure 4.**
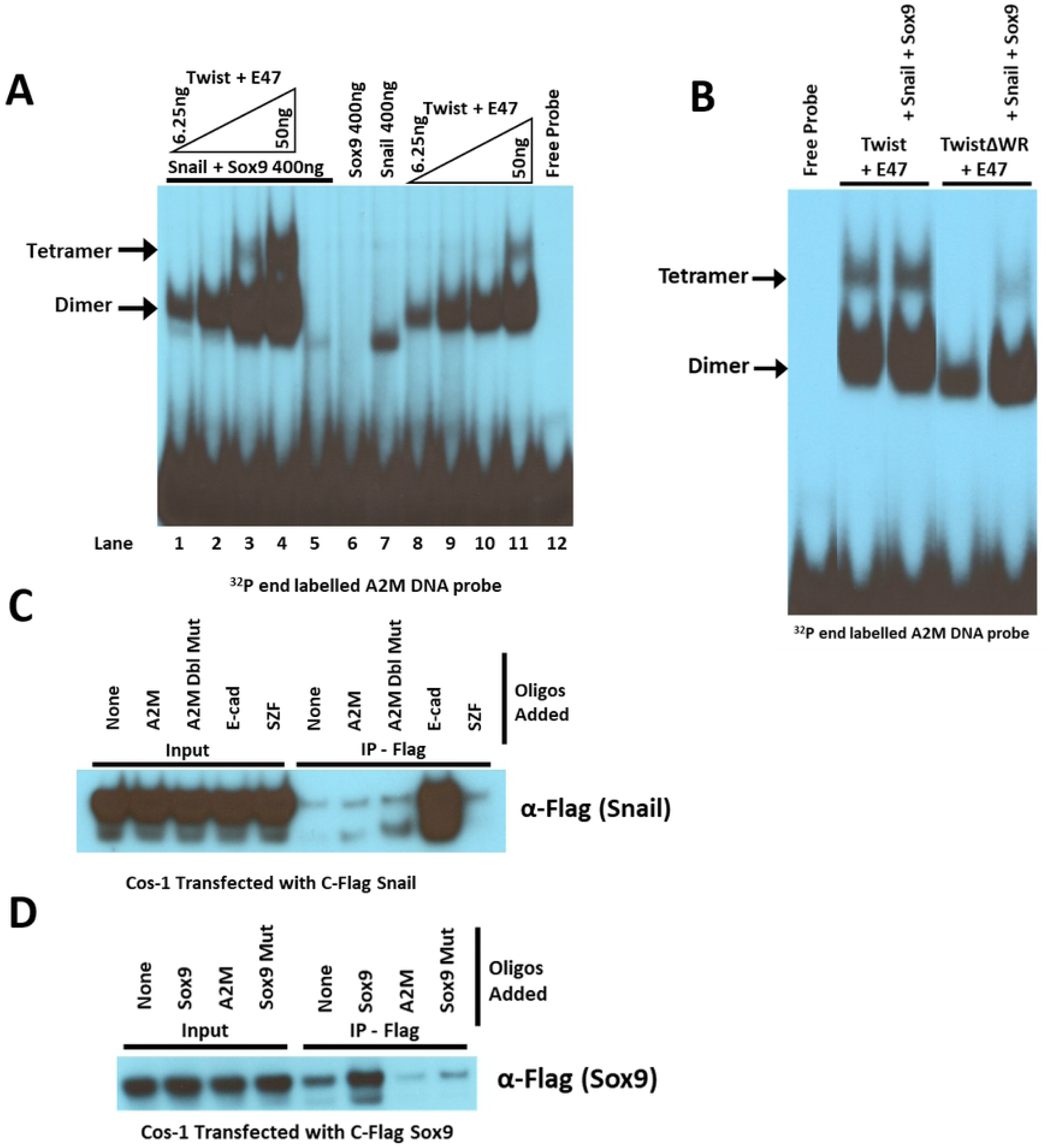
Snail and Sox9 stabilize Twist tetramer formation and are each stabilized by binding to their consensus DNA. A) EMSA with radiolabeled *A2M* DNA probe and titration of recombinant Twist/bHLH+WR-E47/bHLH complex with and without Snail/ZF1-4 and Sox9/DIM+HMG recombinant proteins shows increased binding of higher order complex in presence of Snail and Sox9. B) EMSA with radiolabeled *A2M* DNA probe with recombinant Twist/bHLH+WR-E47/bHLH compared to Twist/bHLHΔWR-E47/bHLH with and without recombinant Snail/ZF1-4 and Sox9/DIM+HMG shows that the presence of Snail and Sox9 are capable of partially restoring the higher order binding which is lost by removal of the Twist WR domain. C) IP(Flag)-Western of transfected Snail-C-Flag from nuclear extracts of Cos-1 cells, with and without addition of *E-cad* DNA binding site and controls shows that Snail is greatly stabilized in presence of its consensus DNA sequence. D) IP(Flag)-Western of transfected Sox9-C-Flag from nuclear extracts of Cos-1 cells, with and without addition of Sox9 dimer DNA binding site DNA binding and controls shows that Sox9 is greatly stabilized in presence of its consensus DNA sequence.

As further evidence of the influence of DNA binding on the folding/conformations of Snail and Sox9, we revisited this for specificity sparked by the data in (Fig 2D) that Snail purification from transfected Cos-1 cells is greatly aided by the addition of the Snail *E-cad* binding site (Fig 2D). We found that this effect is specific for the Snail *E-cad* DNA binding site, as it was not observed using DNA oligos for the *A2M* dyad site or the SZF zinc finger protein consensus binding site (a site unrelated to the E-box) (Fig 4C). A similar effect of DNA binding is also evident upon Sox9 purification (Fig 4D). Thus, the observed increase in soluble protein purification is highly specific to binding their respective binding sites and is highly consistent with the biophysical observations that HMG and Zinc finger domains are quite unstructured prior to DNA binding.

### Reconstitution of the EMTosome

To attempt to reconstruct the EMTosome, we performed a series of triple and quadruple transfections into Cos-1 cells. Overall the data indicate that we can utilize the *A2M* dyad DNA binding site to capture Twist, Snail, and Sox9 proteins from Cos-1 nuclear extracts transfected with Twist, E47, Snail, and Sox9 (Fig 2C). With a triple transfection of Flag-Twist, HA-Snail, and Myc-E47, we observe that both Twist and E47 copurify easily in a DNA affinity procedure using biotinylated *A2M* site (Fig. 5A). We also observe that Snail is capable of copurifying with Twist when Flag-Twist, HA-Snail, and Myc-E47 are triple transfected into cells, but that this copurification is eliminated on addition of the *A2M* DNA oligo, suggesting that complex formation may require both Sox9 and DNA binding for stability. (Fig 5A). When Cos-1 cell extracts containing HA-Twist, Myc-E47, Sox9-C-Flag, and Snail-C-Flag proteins, we observe that Twist and E47 are capable or copurify with Sox9 on a Myc-tag antibody resin (E47). Curiously, under the same conditions HA-Twist is very poorly recovered with HA resin, and no recovery of Snail or Sox9 is seen (Fig 5B). These data suggest that, while we can recover this complex utilizing a DNA binding site of one element of the complex (Fig 2C), the complex cannot be fully recovered in the absence of DNA binding, suggesting that the DNA binding process is not fully separable from complex formation.

**Figure 5.**
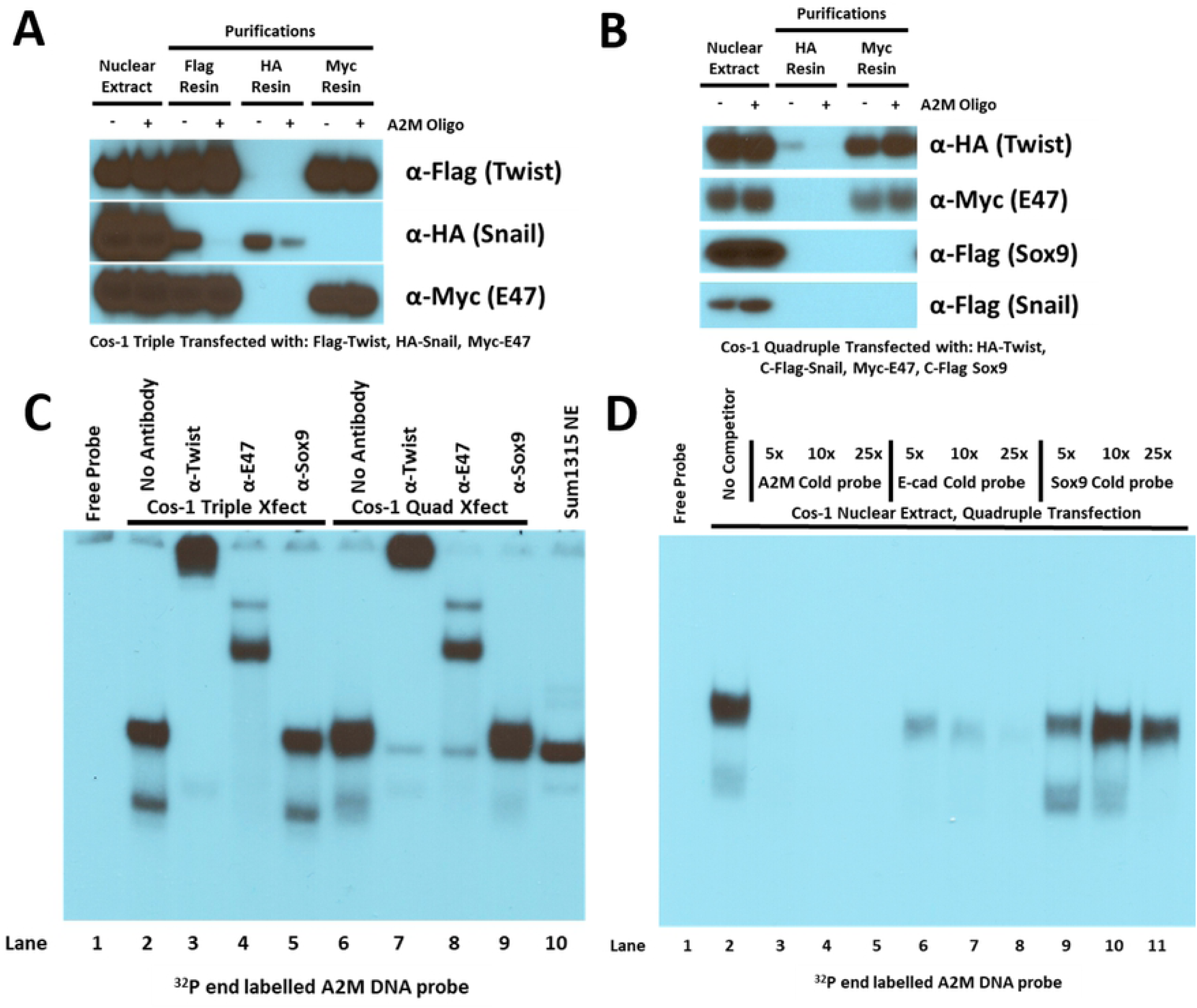
EMTosome can be partially reconstructed with triple and quadruple transfections into COS-1 cells, as shown by Western blot and EMSA antibody shift. A) IP-Western of nuclear extracts from triple transfection of Flag-Twist, HA-Snail, and Myc-E47 into Cos-1 cells shows some interactions between Twist and Snail, and this interaction can be modulated with the Twist dyad E-box DNA. B) IP-Western of nuclear extracts from quadruple transfection of HA-Twist, Myc-E47, Sox9-C-Flag, and Snail-C-Flag into Cos-1 cells show no interactions between Twist-E47 and Snail or Sox9 under these conditions. C) EMSA with radiolabeled A2M DNA probe using antibodies to shift the motility of Cos-1 triple and quadruple transfected nuclear extracts show responsiveness of the complex to Twist and E47 antibodies. D) EMSA with radiolabeled A2M DNA probe using unlabeled DNA competitors A2M, E-cad, and Sox9 dimer as competitors to binding of Cos-1 quadruple transfected nuclear extract shows major inhibition of binding with both A2M and E-cad DNA probes, and slight inhibition of binding with Sox9 consensus DNA.

To further characterize the EMTosome reconstituted in cells, EMSAs with antibody supershifts were done with nuclear extract proteins from quadruple transfected cells (Fig 5C). By EMSA, the major complex co-migrates with the endogenous complex observed many times derived from SUM1315 cells. A marked supershift of this complex is evident when Twist and E47 antibodies are added, but surprisingly there is no effect of Sox9 antibodies. There is a modest but reproducible increase in the abundance of the major complex with the addition of Sox9 to the transfections (Fig 5C, Lane 2 compared to Lane 6), while the mobility of the complex remains overall similar to that of the SUM1315 (Fig 5C, Lane 10). Addition of cold competitor *A2M* and *E-cad* DNA (cold and hot DNAs mixed prior to protein addition) showed that they robustly abrogate binding of the complex to the *A2M* DNA binding site. However, addition of unlabeled Sox9 DNA binding site has a greatly reduced impact on the binding of the complex to the *A2M* DNA binding site (Fig 5D, Lanes 3-5). Altogether, these data suggest that we only partially recapitulate the components and activities of the EMTosome in transfected cells. It’s also likely that additional undefined components and/or post translational modifications are present endogenously in SUM1315 cells which are lacking in transfected Cos-1 cells.

## Discussion

We have taken transcription factor targeted but otherwise unbiased biochemical approach to characterizing the components of a multiprotein complex of Twist, E47, Snail, and Sox9, which are present at high constitutive levels in SUM1315 breast cancer cells that are highly metastatic and display a strong constitutive EMT phenotype. Our main findings can be summarized as follows: 1) Twist protein derived from highly metastatic SUM1315 cells is present in a multiprotein complex which recognizes the dyad E-box with high affinity. 2) This protein complex also contains Snail and Sox9, placing three key TFs for the regulation of EMT into a single complex and this tripartite EMTosome. Moreover, this complex can be reconstituted from cellular proteins derived from transfected cells utilizing the dyad E-box as a ligand. Mutation of one E-box of the dyad Twist *A2M* DNA binding site greatly reduces the ability to bind the endogenous SUM1315 complex, showing high specificity for key DNA binding sites for formation of the holo-complex. 3) When the EMTosome from SUM1315 is challenged by competition with the Snail *E-cad* DNA binding site or the Sox9 DNA binding site, the ability for the complex to bind to the Twist *A2M* binding site is eliminated. Suggesting a mechanism for the regulation of Twist-E47 DNA binding based upon the occupancy of DNA binding sites on the Snail and Sox9 proteins in the complex. Once bound to *A2M* DNA binding site, the EMTosome shows very low dissociation from that site, while Snail and Sox9 DNA targets are capable of still reducing binding, suggesting again a DNA binding based allosteric mechanism for regulation. 4) Individual components of the EMTosome are unstable/poorly folded in the absence of cognate DNA binding showing that formation/folding is facilitated by DNA. Moreover, once formed the components (Snail and Sox9) which do not bind the Twist dyad site directly can still engage their own cognate sites and regulate the overall binding activity of the complex.

Overall, our data support a new working model for TFs implicated in EMT which incorporates important roles for Snail and Sox9 as regulatory components of a large multi-protein complex on an EMT target gene, but which do not see the target gene directly. (Fig 6). Normally, Twist binds to a dyad E-box, along with the rest of the EMTosome, where it regulates the transcription of the associated gene (Fig 6A). But when Snail or SOX9 bind to their DNA consensus sequences, recruiting the EMTosome to those sequences, they restrict the transcriptional regulation of Twist by eliminating the DNA binding capabilities (Fig 6, B and C). We suspect that WR domain is only properly available for binding to Snail and Sox9 when Twist-E47 tetramer is bound to its dyad site. In this scenario, a DNA bound WR-domain is a signal that Twist target genes are “on” and possibly, by tying up Snail in the EMTosome, this keeps direct Snail targets off or not yet regulated. The ability of Snail in the EMTosome to engage its high affinity site, leading to instability or abolishment of Twist-E47 binding to its target dyad site is then a toggle key switch between Twist regulated target genes and Snail regulated target genes. This would further differentiate Twist-E47 target genes from direct Snail target genes more so than just the differences in the nucleotide sequence of the E-boxes do in the E-cads site and the dyad sites. The dyad symmetric E-box clearly does this itself as 70% of sites obtained in our initial Chip-seq experiments (15) were dyad symmetric sites with a 5-base spacing. Moreover, most of these dyad E-box sites show little relative affinity for Snail. It is an interesting mechanism by which sequentially regulated target genes in a genetic program are regulated by apparent allosteric regulation of DNA binding domains of proteins present in the same macromolecular complex. It is also noteworthy that Twist is largely defined as an activator of transcription whereas Snail is largely a repressor: this further supports both a physical and functional separation of regulatory sub-programs in the EMT phenotype.

**Figure 6.**
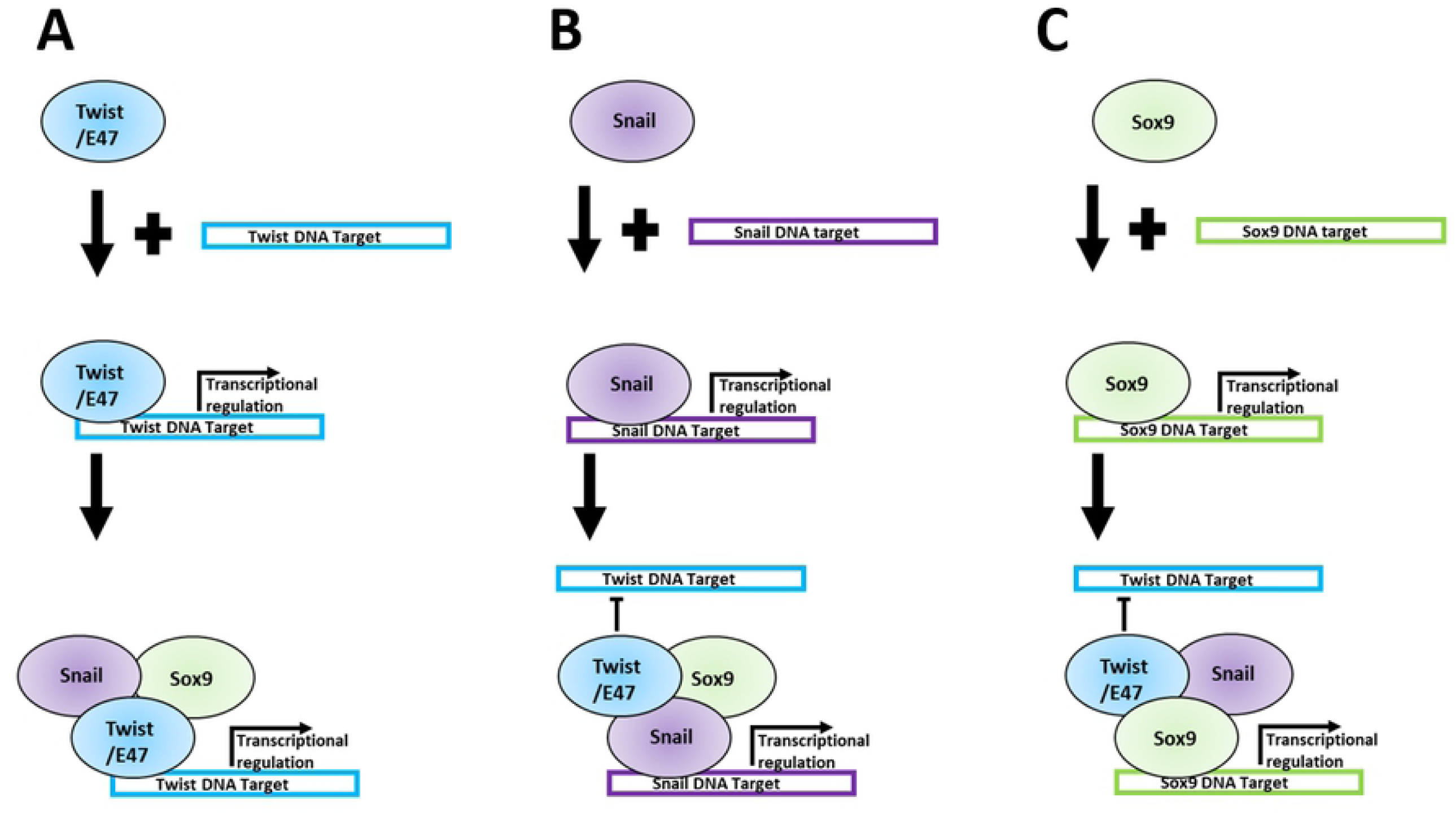
Model of the Molecular Mechanisms of the EMTosome. A) Twist/E47 bind to dyad E-box target DNA, and form the EMTosome complex on this DNA, resulting in transcriptional regulation of Twist target genes. B) Snail binds to single E-box target DNA, and forms the EMTosome on that target DNA, but by doing so allosterically eliminates the ability for Twist/E47 to bind to their target DNA, preventing transcriptional regulation by those factors. C) Sox9 binds to target DNA, and forms the EMTosome on that target DNA, but like Snail, eliminates the ability for Twist/E47 to bind to their target DNA, preventing transcriptional regulation by those factors.

In previous studies, the interactions and regulation between Twist and Sox9 for chondrogenesis have already been categorized according to their genetic regulation of each other, rather than by biochemical and molecular interactions (13). Most relevant are the number of studies showing a strong genetic interaction between Twist, Snail and Sox9 in a variety of forward and reverse genetically manipulatable systems in both embryonic development and cancer models. Our discoveries reported here greatly complement and extend these by showing high affinity protein-protein interactions amongst these proteins while also proposing a novel mode of their cross regulation.

Other studies showed a repressive effect of Twist on RunX proteins and osteogenesis (20), a regulatory effect of Sox9 on Snail stabilization (14), and the impact of downregulated Sox9 on many traits of metastatic cancer such as invasiveness, proliferation, and the EMT phenotype (11). This gives us a clear picture of the large role that these TFs are playing in the chondrogenesis versus osteogenesis portion of cellular differentiation, as well as maintenance of a stem-like phenotype. Combined with the more EMT specific roles of these TFs (21), such as direct interactions of Twist with Snail (22), we can see how these similar but distinct genetic processes can contribute to metastatic spread, and understanding the mechanistic interactions between these disparate TFs is crucial for designing targeted therapeutic approaches (23).

Recent research supports the premise that partial EMT, in which cells display both epithelial and mesenchymal traits simultaneously, may be relevant to cancer metastasis (24-26), suggesting that further research into the specific biochemical mechanisms of this partial EMT phenotype are necessary. Our finding that several EMT TFs are sequestered into a single complex, the EMTosome, and the occupancy of DNA binding sites by one member impacts the DNA binding ability/affinity of the other complex members, may offer a mechanistic explanation for the partial EMT state phenotype. Our model proposes an efficient mechanism for controlling specific genetic differentiation programs. By having different factors in a single multi-protein complex, different DNA binding interfaces are available to be recruited at different target genes, allowing for incomplete EMT phenotypes, such as loss of partial cellular junctions with added motility, allowing cell clusters to metastasize rather than individual cells (25). In addition, other mutations in this complex, such as the WR domain of Twist, have been shown to contribute to other differentiation based pathologies, such as Saethre-Chozen Syndrome and Craniosyntosis (27, 28), where haploinsufficiency results in defects in the differentiation of the neural crest (29-31).

## Conclusions

In summary, multiple EMT TFs, Twist-E47, Snail, and Sox9, are sequestered into a single multiprotein complex, we have designated the EMTosome. The DNA binding activities of these TFs appears to cause allosteric changes which can result in increased stability and modulate the DNA binding activities of other TFs in the complex. This provides a mechanism for the hierarchal, sequential series of genetic regulatory programs which must activate and deactivate in a precise way to drive EMT. DNA binding of one protein and then recruitment of the other proteins in the EMTosome away from other target DNA also provides a biochemical and molecular mechanism by which partial EMT, a potent driver of cancer metastasis, can occur.

## Acknowledgements

The authors would like to thank The Wistar Institute Core Facilities, especially the proteomics and genomics core facilities and Perelman School of Medicine’s Biochemistry and Molecular Biophysics Graduate Program for all their support. Thanks to Maureen Murphy at The Wistar Institute for internally reviewing the manuscript.

---

**Supporting information Figure 1.**
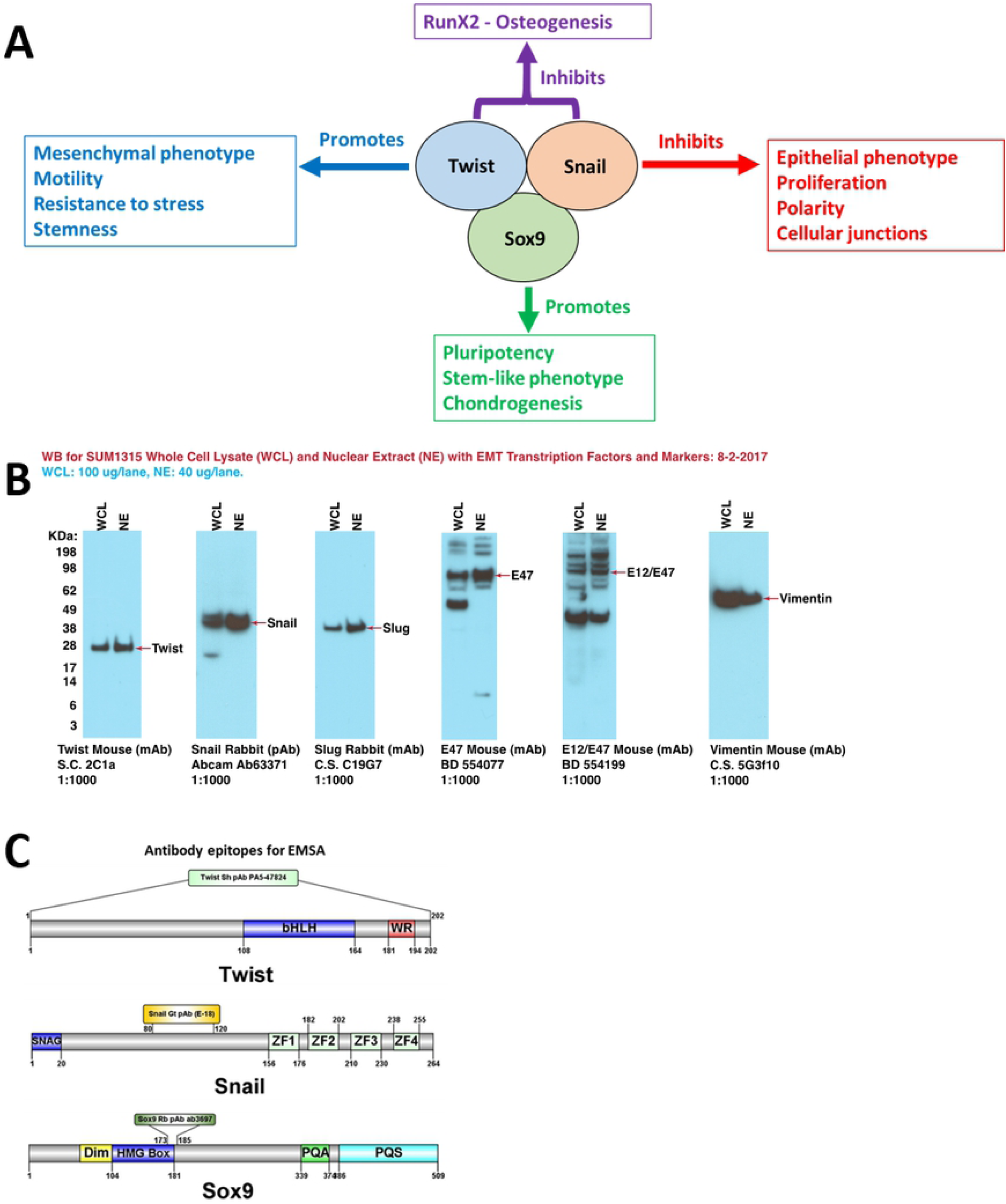
A) Diagram of some of the processes being regulated by the members of the EMTosome protein complex, highlighting the many similar and related processes regulated by different members of this complex. B) Sum1315 metastatic breast cancer cells show markers for the EMT phenotype, as shown by Western blots of both whole cell lysate and nuclear extract from Sum1315 cells. C) Epitope maps of the antibodies used in EMSA to identify EMTosome (Fig 1D)

**Supporting information Figure 2.**
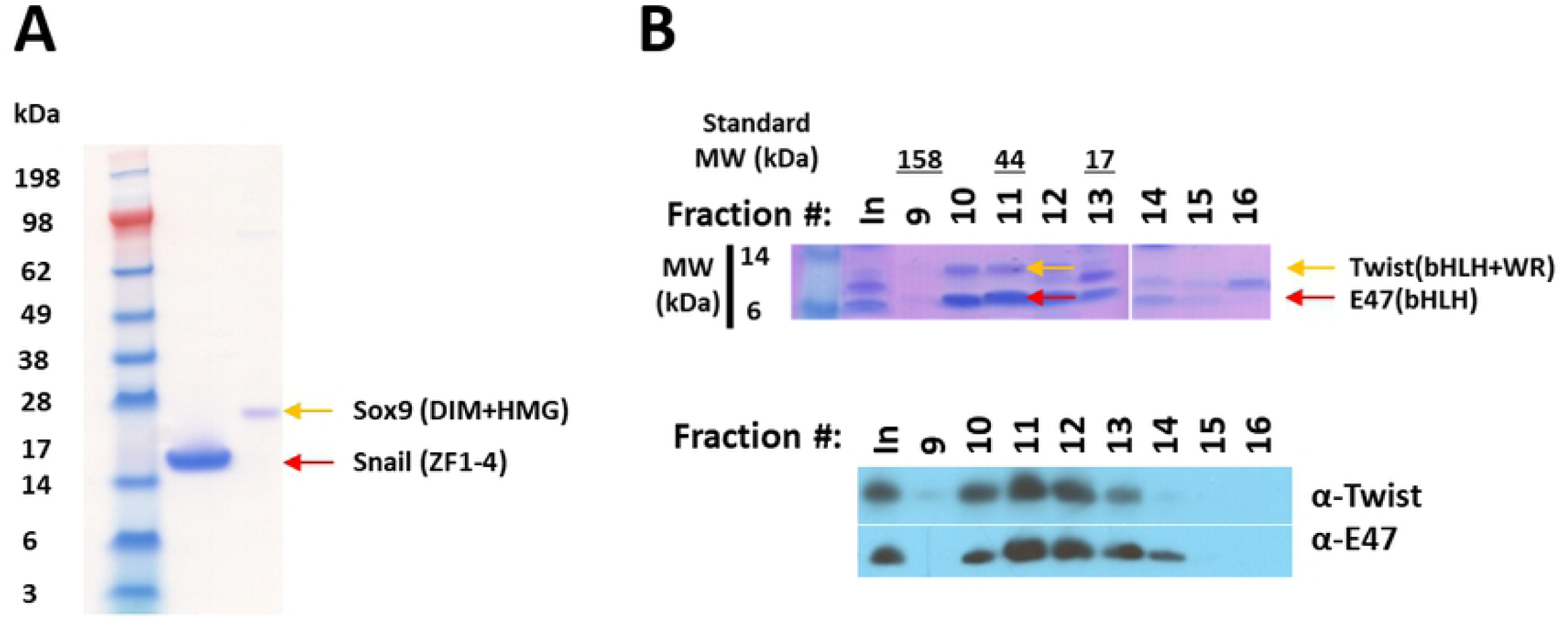
Purification of recombinant constructs used show clean well separated proteins. A) Denaturing purification of His-Snail/ZF1-4 and His-Sox9/DIM+HMG utilizing nickel-NTA beads and then SDS-PAGE with coomassie staining. B) SDS-PAGE of fractions from Size Exclusion Chromatography after native co-expression and copurification of Twist/bHLH+WR and E47/bHLH from cobalt beads with both coomassie staining and Western Blot.

